# Reactive oxygen species mediate thylakoid membrane remodeling and triacylglycerol synthesis under nitrogen starvation in the alga *Chlorella sorokiniana*

**DOI:** 10.1101/2021.05.15.444036

**Authors:** Jithesh Vijayan, Nishikant Wase, Kan Liu, Chi Zhang, Wayne R. Riekhof

**Affiliations:** School of Biological Sciences, University of Nebraska-Lincoln; Department of Biochemistry, University of Nebraska-Lincoln; Center for Plant Science Innovation, University of Nebraska-Lincoln; School of Medicine, University of Virginia System Charlottesville, VA

## Abstract

Many microbes accumulate energy storage molecules such as triglycerides and starch during nutrient limitation. In eukaryotic green algae grown under nitrogen limiting conditions, triglyceride accumulation is coupled with chlorosis and growth arrest. In this study we show that accumulation of reactive oxygen species (ROS) under nitrogen limitation in the microalga *Chlorella sorokiniana* is involved in thylakoid membrane remodeling, leading to chlorosis. We show that ROS accumulation under nitrogen limitation is an active process involving downregulation of expression of ROS-quenching enzymes, such as superoxide dismutases, catalase, peroxiredoxin, and glutathione peroxidase-like, and upregulation of enzymes involved in generating ROS, such as NADPH oxidase, xanthine oxidase and amine oxidases. Expression of enzymes involved in ascorbate and glutathione metabolism are also affected under these conditions. Quenching ROS under nitrogen limitation reduces TAG accumulation, adding additional evidence for the role of ROS signaling in the process.

## INTRODUCTION

Microalgae are considered a source of renewable biofuels as they are fast growing and do not compete with crops for arable land (Hu et al. 2008). Significant investment has been made in the past two decades toward developing algae as a reliable biofuel and bioproduct feedstock, while coupling biomass production to waste remediation (Allen et al. 2018). Microalgae, like many other microbes, accumulate storage molecules when experiencing unfavorable growth conditions (Zhu et al., 2016). Starch and oil are products of interest as sources of biofuel. Under nutrient limitation conditions, microalgae like *Chlorella* spp., *Chlamydomonas reinhardtii*, and *Nanochloropsis* spp. accumulate oil (Wase et al. 2014; Miller et al. 2010; Stephenson et al. 2010; X. N. Ma et al. 2016; Jiang et al. 2019). A great deal of effort has gone into understanding the mechanisms of metabolic remodeling and regulation of oil accumulation under nitrogen starvation (Miller et al. 2010; Wase et al. 2017; 2019). Many different enzymes involved in the process of channeling photosynthate to oil have been identified (Boyle et al. 2012; Shtaida et al. 2014). Nitrogen limitation is a strong inducer of oil accumulation in different microalgae, but invariably growth is reduced and carbon fixation is severely compromised. This is an unfavorable factor when considering the industrial use of algae as a feedstock for waste-remediation and biofuel production. Furthermore, growth arrest and accumulation of oil is accompanied by the turnover of chlorophyll and chlorosis, also known as the “degreening” of cells.

Under nitrogen limiting conditions, fatty acids for synthesis of triacylglycerol (TAG) are derived from both *de novo* synthesis and by remodeling of glycerolipids from preexisting membranes. Various enzymes involved in membrane remodeling have been reported in *Chlamydomonas reinhardtii* (Li, et al., 2012a; Li, et al., 2012b; Yoon, et al., 2012) which act on different glycerolipids in membranes to release fatty acid for TAG synthesis. Upon nitrogen limitation, microalgae like *Chlamydomonas reinhardtii* degrade monogalactosyldiacylglycerol (MGDG), a plastidial membrane lipid, to synthesize triacylglycerol (TAG) (Li, et al., 2012).

All aerobic organisms use oxygen as an electron acceptor in respiration. Various pathways, such as mitochondrial electron transport, nitric oxide synthase, and peroxisomal β-oxidation lead to production of reactive oxygen species (ROS) (Møller 2001). Different forms of ROS are present in cells, such as superoxide (O_2_^-^), hydroxyl radical (. OH-), singlet oxygen and hydrogen peroxide (H_2_O_2_). Each of these reactive species have different half-lives and chemical properties which affects the biochemistry and physiology of cells (Foyer and Noctor 2009).

Cellular redox and ROS homeostasis is carefully balanced by an elaborate antioxidant defense system which includes enzymes such as superoxide dismutase (SOD), catalase, glutathione peroxidases (GPX), ascorbate peroxidases (APX) and low molecular weight scavengers such as uric acid, ascorbate, glutathione, carotenoids and flavonoids (Czarnocka and Karpiński 2018). Studies over the past two decades have revealed ROS as a fine-tuned signaling system that elicits both localized and global responses. Under different biotic and abiotic stress conditions, ROS signaling has been implicated to play a key role in stress responses and acclimation (Ben Rejeb et al. 2015; Gupta, Palma, and Corpas 2015; He et al. 2017) which is prevalent in both animals and plants (Foyer and Noctor 2009; Schieber and Chandel 2014).

In this study we provide evidence that ROS signaling is involved in the nitrogen starvation response in *Chlorella sorokiniana*, an industrial microalga of the family *Chlorophyceae*. We show that *C. sorokiniana* uses transcriptional and post-translational regulatory mechanisms to accumulate ROS, and that ROS signaling mediates chloroplast membrane remodeling leading to chlorosis and oil accumulation.

## RESULTS

### Nitrogen limitation causes growth arrest coupled with oil accumulation

Nitrogen is an essential macronutrient for the growth of all organisms, being necessary for synthesis of nucleic acids, amino acids, and the amino-acid derived head groups of certain lipids like phosphatidylcholine and phosphatidylethanolamine. In *Chlorella sorokiniana*, nitrogen limitation leads to growth arrest, as shown in Fig. 1A. We observe a decrease in growth as early as 9 hours after inoculation into nitrogen-deficient acetate-containing media. *C. sorokiniana*, like many other microalgae, accumulate oil droplets when deprived of nitrogen. Fig. 1B shows Nile Red stained *C. sorokiniana* cells under nitrogen-limited and replete conditions, with lipid droplet-bound Nile Red fluorescence evident as green puncta.

**Fig 1.**
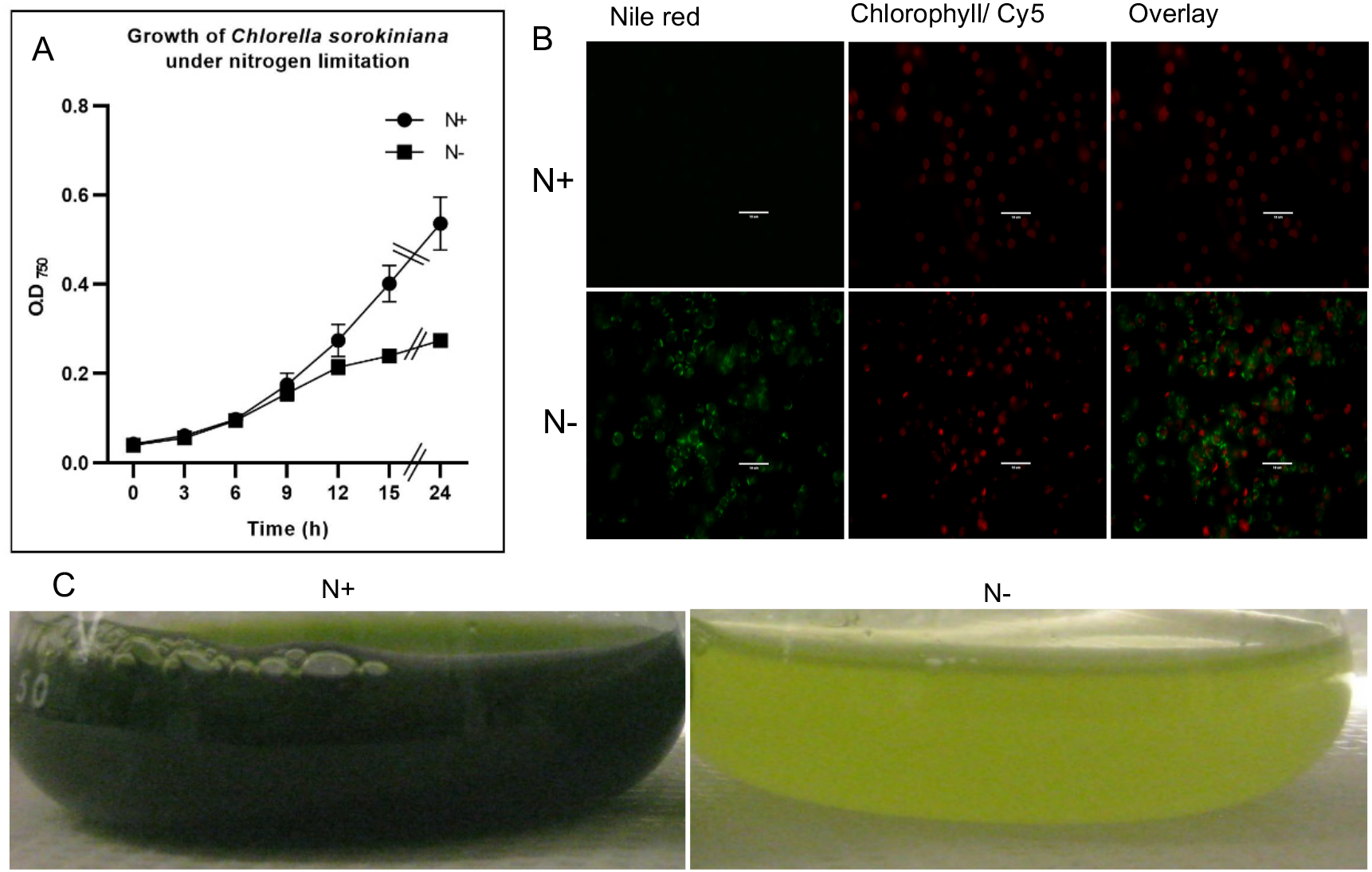
Growth and oil accumulation of *Chlorella sorokiniana* under nitrogen limitation. A) Growth curves of *Chlorella sorokiniana* in N-replete (N+, 7.5 mM) and N-limited media (N-, 0.25 mM). Cultures were inoculated with 107 cells/ml and the mean +/- SD of three biological replicates is presented. B) Lipid droplet accumulation under N-limitation. N+ and N-cultures were prepared as for the growth curves. Cultures were grown for 48 h and collected, stained with Nile Red to visualize lipid droplets, and imaged by fluorescence microscopy. C) Culture of *Chlorella sorokiniana* inoculated with 107 cells/ml as viewed after 48 hrs.

### Chloroplast membrane remodeling accompanies chlorosis

Nitrogen limitation in *C. sorokiania* causes severe chlorosis as depicted in Fig. 1C. Chlorosis is caused by degeneration of chloroplast membranes, accompanied by chlorophyll degradation. We carried out transmission electron microscopy (TEM) to observe the effects of nitrogen limitation on membrane integrity of chloroplasts. Fig. 2A shows the TEM micrographs of nitrogen limited cells. Dense thylakoid membranes in control cultures (+N) are significantly reduced after 4 days of nitrogen starvation (-N). One of the characteristics of nitrogen limitation in microalgal cells is the substantial reduction in the levels of monogalactosyldiacylglycerol (MGDG);(Li, et al., 2012). Membrane lipids of nitrogen-replete cultures were labeled to saturation with 1,2-^14^C-acetate, and during a chase period upon removal of labeled acetate and induction of N-limitation, we see the degradation of MGDG as shown in Figs 2B and 2C. Other lipids, such as phosphatidylcholine and phosphatylethanolamine, are also degraded. Fatty acids released from these lipids are channeled into TAG as seen by the increase in radiolabeled TAG during the N-limited chase period of growth (Fig. 2B).

**Fig 2.**
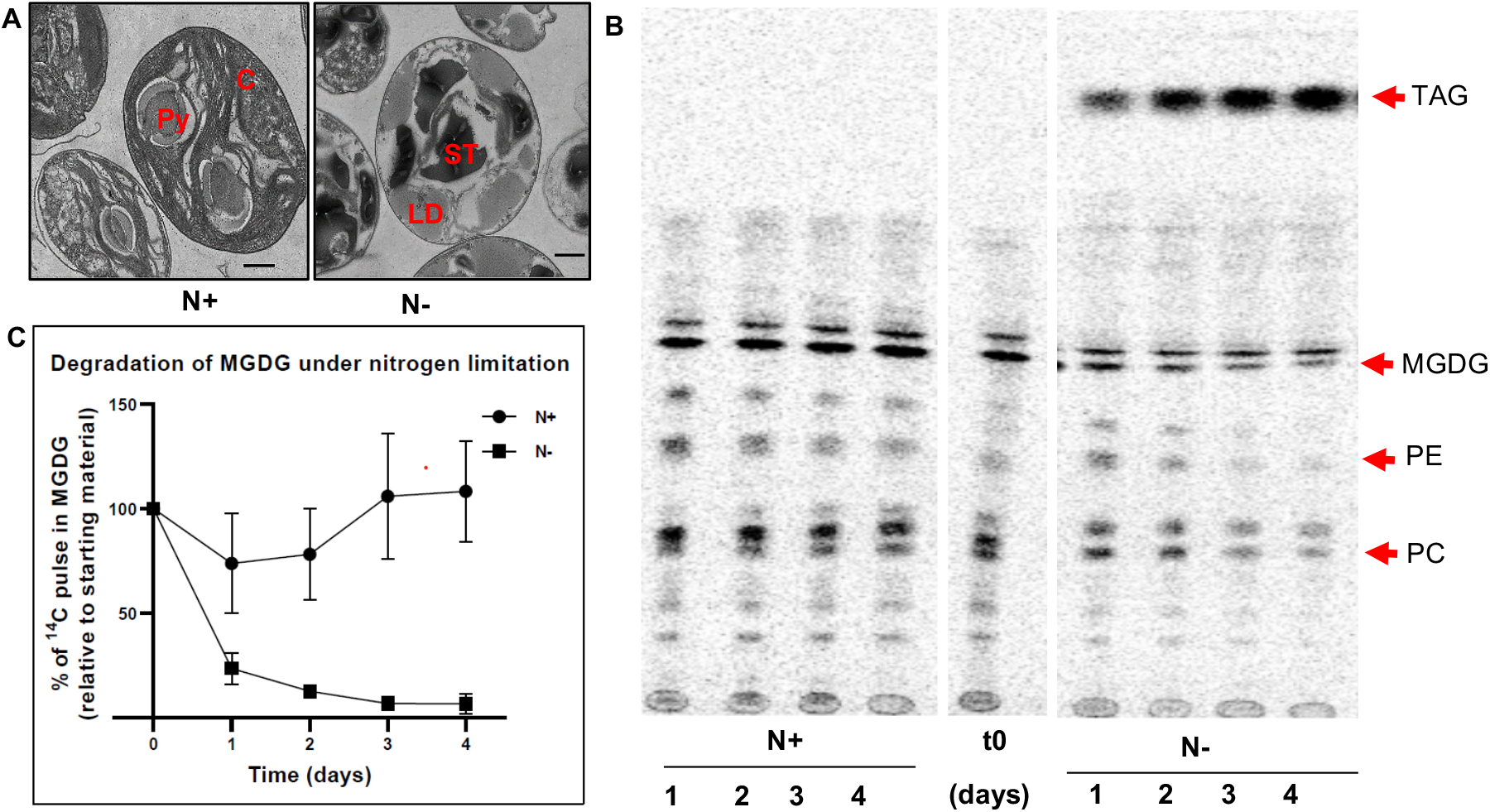
Chloroplast membrane remodeling under nitrogen limitation. A) TEM image of *Chlorella sorokiniana* cells under N+ or N-conditions 4 days post inoculation. C: Chloroplast, PY: Pyrenoid body, ST: Starch granule, LD: Lipid Droplet. Scale bars indicate 800 nm. B) Representative TLC-radiograph of cells prelabeled with ^14^C - acetate and then grown in N+ or N-media, indicating the degradation of preformed lipids under N-conditions. TAG, Triacylglycerol; MGDG, Monogalactosyldiacylglycerol; PE, phosphatylethanolamine; PC, phosphatidylcholine. C) Quantitative measurement of ^14^C-containing MGDG degradation upon N-limitation. Measurements are relative to the amount to ^14^C pulse present in MGDG at the initial time point (t0).

### Transcriptomic analysis of nitrogen limited cells

To further understand the mechanisms underlying chloroplast membrane degradation and oil accumulation under nitrogen limitation, we carried out RNAseq analysis. We chose 9 and 24 hours after inoculation for sampling as early and late time points respectively. Nine hours was chosen as the early time point for sampling, as this was the time at which a decline in growth rate in N-limited culture is initially observed relative to an N-replete culture (Fig. 1A). We find that 16 % of the genes are significantly differentially expressed in the early time point at 9 hours and 22 % show altered expression at the 24 hour time point (Table 1). This shows that nitrogen starvation response starts earlier than 9 hours, i.e. before a significant reduction in growth is noticeable.

**Table 1:** Account of significantly changed genes.

| Condition | N vs T-9h (%) | N vs T-24h (%) |
| --- | --- | --- |
| Down | 1076 (6.99 %) | 1693 (11%) |
| No significant change | 12805 | 11895 |
| Up | 1501 (9.76%) | 1794 (11.66%) |
| Total | 15382 | 15382 |

**Number of genes whose expression were significantly changed.** Adjusted p-value <0.05 and log2 fold change of 2. Percent values in parenthesis indicate the percentage relative to total number of genes in the genome.

### A transcriptomic signature of ROS accumulation under nitrogen limitation

From the RNAseq data, we find that many genes encoding enzymes involved in quenching ROS were downregulated. Enzymes such as superoxide dismutase (SOD), catalase, and glutathione peroxidases quench and detoxify ROS that are generated during various cellular process such as photosynthesis and mitochondrial electron transport (Asada 2006; Møller 2001). Expression of superoxide dismutases (SOD1, SOD2, SOD3) and catalase are downregulated as shown in Fig.3a. SOD reduces superoxide radicals to H_2_O_2_ which is further reduced by a variety of enzymes such as peroxidases and catalase (Czarnocka and Karpiński 2018). Expression of genes involved in ascorbate (AsA) metabolism are also significantly altered (Fig. 3b). We see that genes involved in synthesis of AsA from GDP-mannose such as GDP-d-mannose pyrophosphorylase (GMnPp), GDP-mannose-3,5-epimerase 1 (GMnEpi), GDP-L-Galactose Phosphorylase 1 (GGlcPp), L-Galactose-1-Phosphate Phosphatase (GlcPiPtase) and L-Galactono-1,4-Lactone Dehydrogenase (GlcLacD) (Smirnoff 2018) are downregulated indicating that the biosynthesis of AsA is decreased under N-limitation. While the expression of AsA synthesis genes are downregulated, AsA reducing enzymes such as monodehydroascorbate reductase (MDHAR) and dehydroascorbate reductase (DHAR) are upregulated (Fig. 3B). This indicates that the availability of AsA may be regulated by regenerating the oxidized MDHA or DHA. Glutathione reductase (GR) which regenerates GSH from oxidized form of GSSG using NADH as a substrate (Foyer and Noctor 2009) is also downregulated (Fig. 3C). Expression of glutathione peroxidase-like enzymes (GPrxL-1 and 2) which possibly use GSH to reduce hydrogen peroxide are also downregulated. This indicates that GSH mediated quenching of H_2_O_2_ could be decreased as well.

**Fig 3.**
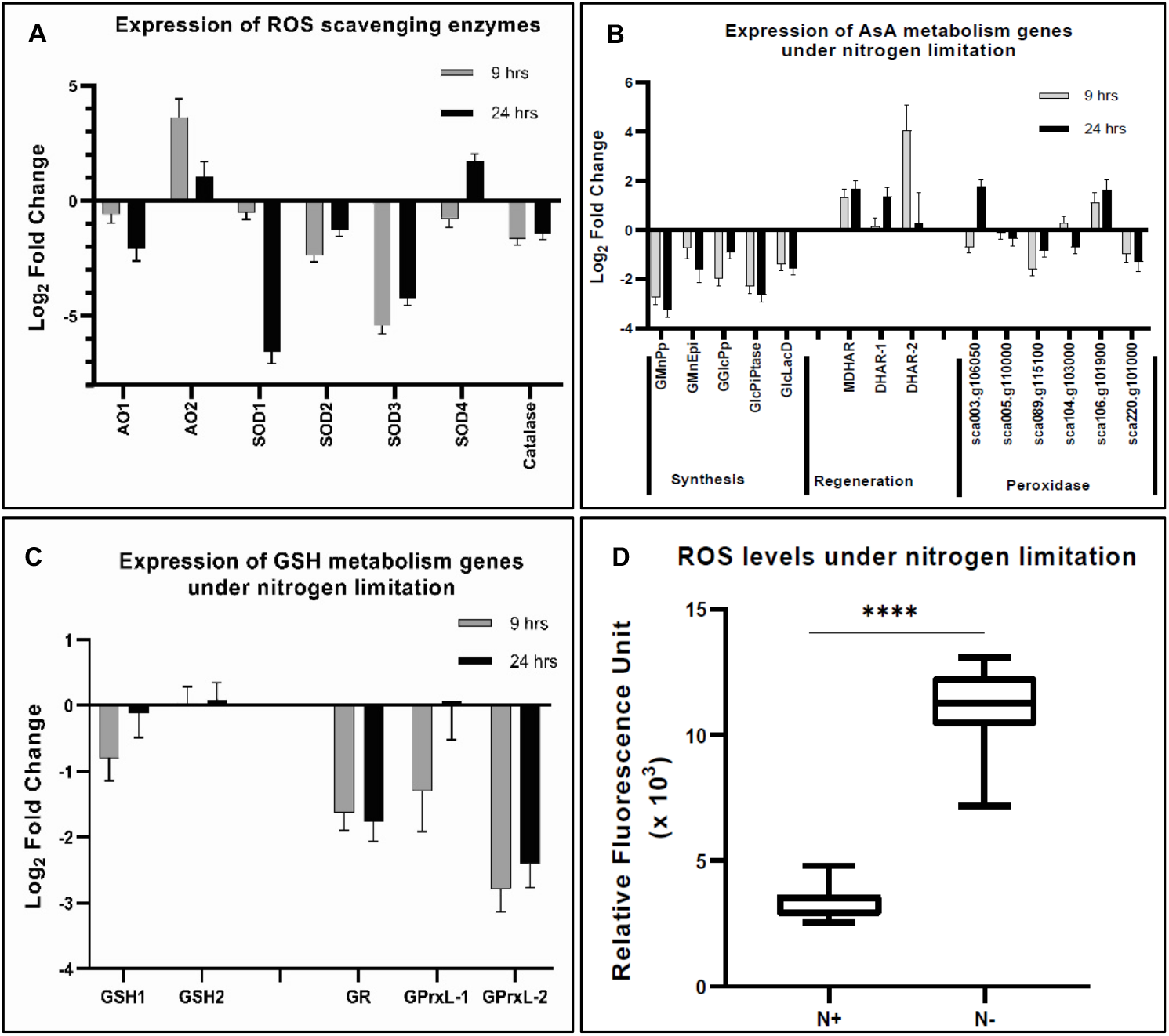
Reactive oxygen species under nitrogen limitation. A) Expression of enzymes involved scavenging ROS. AO: Alternative oxidase, SOD: Superoxide Dismutase. B) Expression of genes involved in ascorbate (AsA) metabolism. Genes are classified as those involved in AsA synthesis, regeneration and AsA dependent peroxidases. GMnPp: GDP-d-Mannose Pyrophosphorylase, GMnEpi: GDP-Mannose 3,5-pimerase 1, GGlcPp: GDP-L-Galactose Phosphorylase 1, GlcPiPtase: L-Galactose-1-Phosphate Phosphatase, GlcLacD: L-Galactono-1,4-Lactone Dehydrogenase, MDHAR: Monodehydroascorbate Reductase, and DHAR: Dehydroascorbate Reductase. C) Expression of genes involved in glutathione metabolism. GSH1: Gamma-glutamylcysteine Synthetase and GSH2: Glutathione Synthetase are involved in glutathione synthesis. GR: glutathione reductase regenerates reduced glutathione (GSH) from oxidized form of glutathione (GSSG). GPrx : glutathione peroxidases oxidizes GSH to quench ROS and form GSSG. D) ROS level as measured by DCFDA fluorescence under N-lim condition after 24 hours of inoculation in appropriate media. The values are normalized to optical density. Error bars indicate SD (n=3).

Downregulation of the expression of these key enzymes involved in quenching ROS, as shown in Figs. 3A-C, may indicate that under N-limitation, *C. sorokiniana* cells accumulate ROS in a transcriptionally regulated manner. To test this hypothesis, we measured intracellular ROS levels in N-limited cells using a cell-permeable ROS-sensitive fluorescent dye, 2’-7’- dichlorodihydrofluorescein diacetate (DCFDA). We see that ROS levels in N-limited *C. sorokianiana* cells are significantly higher than that of the control cells (Fig. 3D).

To gain a more complete picture of the redox regulation under N-limitation, we tabulated the expression profile of various proteins of the ferredoxin, peroxiredoxin, thioredoxin, and glutaredoxin families. There are significant changes in the expression of many ferredoxin proteins that we examined (Fig. 4a). These proteins can function as a reservoir of redox potential, accepting or donating electrons to various biochemical processes including photosynthesis (Schürmann and Buchanan 2008). Expression of four peroxiredoxin proteins are significantly downregulated (Fig. 4b). Along with catalase and peroxidases, these proteins are involved in scavenging H_2_O_2_ (Rhee and Woo 2011). Downregulation of peroxiredoxin proteins indicates that yet another mechanism of ROS scavenging could be perturbed in the cells under N-limitation. Expression of many thioredoxin proteins are also changed upon N-limitation (Fig. 4c). Thioredoxin proteins are known to be involved in signaling by modification of cysteine residues of proteins (Baumann and Juttner 2002). We also see that thioredoxin reducing enzymes such as ferredoxin-thioredoxin reductases 1 and 2 (Frdxn TrxRed1 and 2) (Schürmann and Buchanan 2008) are downregulated (Fig. 4D) indicating that thioredoxin mediated redox signaling is changed under N-limitation. Glutaredoxins, which share functional similarity to thioredoxins (Meyer et al. 2008), are also differentially regulated as shown in Fig. 4E. These proteins were identified based on sequence homology to characterized proteins of the class and hence needs further characterization to verify/ validate the function.

**Fig 4:**
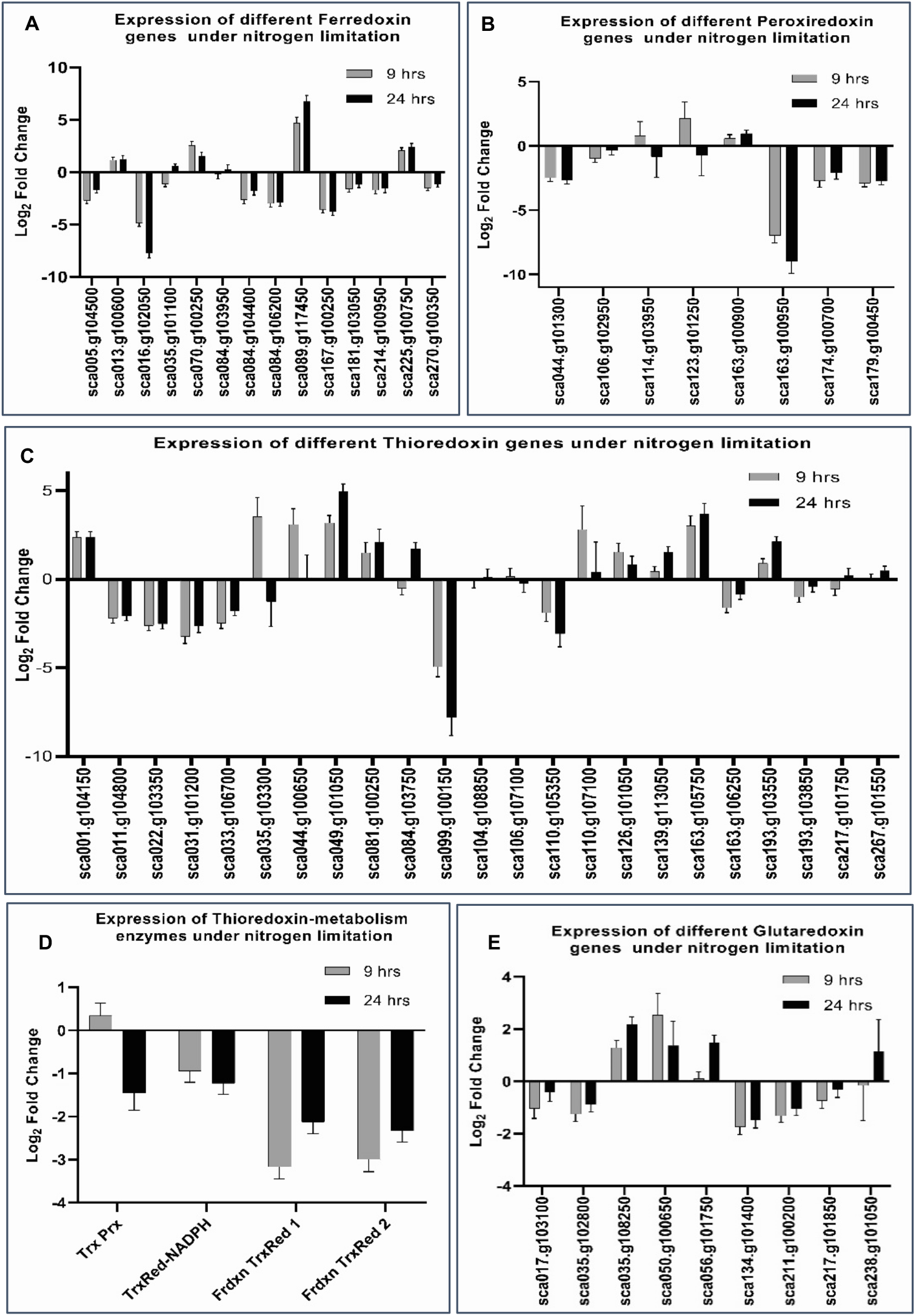
Expression of redoxin proteins during nitrogen limitation. Expression of different Ferredoxin (A), Peroxiredoxin (B), Thioredoxin-like (C), Glutaredoxin-like (F) proteins during nitrogen limitation. D) Expression of Thioredoxin-metabolism enzymes under nitrogen limitation. Trx Prx: Thioredoxin peroxidase, TrxRed-NADPH: NADPH dependent Thioredoxin reductase, Frdxn TrxRed1 and 2: Ferredoxin Thioredoxin reductase1 and 2. E) Expression of different glutaredoxins under nitrogen limitation.

Given the fact that SODs, catalase, peroxidases, ferredoxins and peroxiredoxins are downregulated, we propose that ROS accumulation in N-limited *C. sorokiniana* cells is mediated, at least partially, by down-regulation of ROS scavenging enzymes leading to a more oxidative cellular environment.

### ROS and chloroplast remodeling

We next asked if the observed increase in ROS level induces the degradation of MGDG and other membrane remodeling events. To test this hypothesis, we treated cells grown under N- replete conditions with diethyldithiocarbamate (DDC), a known inhibitor of superoxide dismutase (Heikkila, et al., 1976). Treatment of cells with 250 μM DDC resulted in degradation of MGDG, as shown in Fig 5A, similar to N-limited cells. MGDG degradation in DDC treated cells is not as extensive as in nitrogen limited cells, possibly because DDC inhibits SOD, which catalyzes the conversion of superoxide (O_2_^-^) to hydrogen peroxide (H_2_O_2_). This process can also occur spontaneously at a slower rate, and other ROS quenching mechanisms are active as well, which may decrease the accumulation of ROS relative to that found in N-limited cells. Conversely, we wanted to know if preventing accumulation of ROS under nitrogen limitation can prevent membrane remodeling. To test this hypothesis, we used ascorbate to quench cellular hydrogen peroxide (Smirnoff 2018). We measured ROS levels in N-limited cells treated with and without 500 μM ascorbate. We found that the ROS levels were lower in ascorbate treated cells at 24 hours after onset of N-limitation (Fig. 5B). This reduction in ROS under nitrogen limitation is coupled with a decrease in MGDG degradation as measured after 6 hours of N-limitation (Fig. 5C). We also notice that when cells are treated with ascorbate under N-limitation, TAG accumulation is reduced, as seen in Fig. 5D. Taken together, these results indicate that accumulation of ROS under N-limitation plays a vital role in chloroplast membrane remodeling, by initiating the degradation of MGDG and channeling FA into TAG accumulation.

**Fig 5.**
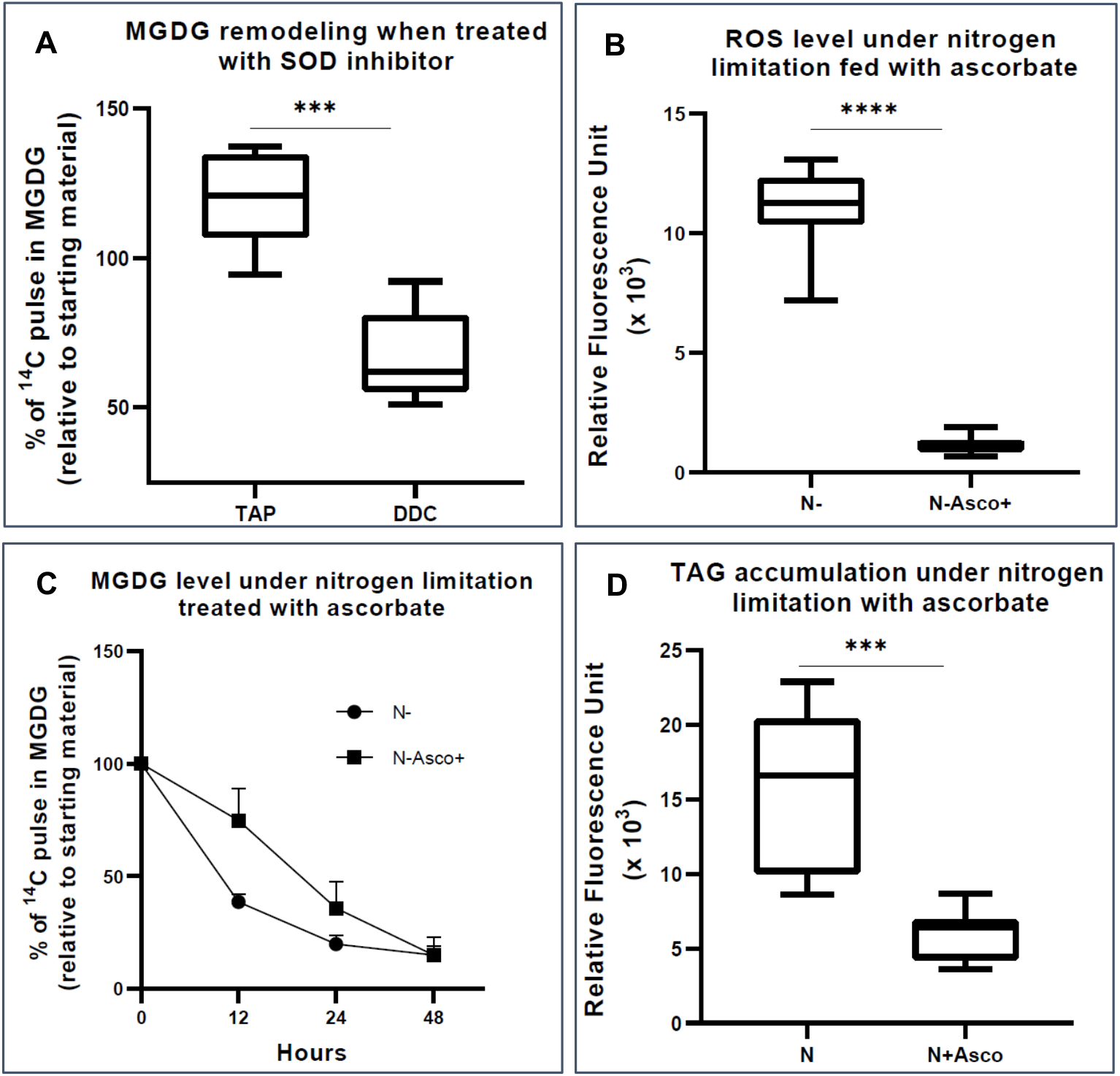
Quenching of ROS under nitrogen limitation and induction of ROS under nitrogen replete conditions. A) Quantitative measurement of MGDG degradation in Nitrogen replete media (TAP) when treated with 250 uM Diethyldithiocarbomate (DDC) after 24 hrs. B) ROS level under nitrogen limited media fed with Ascorbate, as measured by DCFDA fluorescence after 24 hours of inoculation. Fluorescence values are normalized to O.D. C) Quantitative measurement of MGDG degradation under N *lim* media when fed with 500 μM Ascorbate. D) Level of TAG accumulation as measured by nile red, under nitrogen limitation when fed with Ascorbate

### Sources of ROS under N-limitation

Photosynthesis is a prominent source of superoxide in all photosynthetic organisms (Czarnocka and Karpiński 2018). We tested the hypothesis that does the increase in ROS observed in N-limited cells is derived from photosynthesis, and ROS accumulation mediated by downregulation of ROS quenching enzymes. To decipher this, we measured ROS levels in cells grown under continuous light or dark in N-limited conditions. We predicted that ROS levels would be significantly lower in N-limited cells when grown in dark. However, we found that there was no appreciable reduction in ROS levels in dark-grown cells, as shown in Fig 6A. This is also reflected in the lipid profile, as we did not notice any appreciable difference in degradation of MGDG as shown in the TLC autoradiogram (Fig. 6B) and the quantitative measurement of MGDG uniformly labeled with 1,2-^14^C-acetate (Fig. 6C). This indicates that the ROS associated with N-limited lipid remodeling in *C. sorokiniana* comes from sources other than photosynthesis.

**Fig 6.**
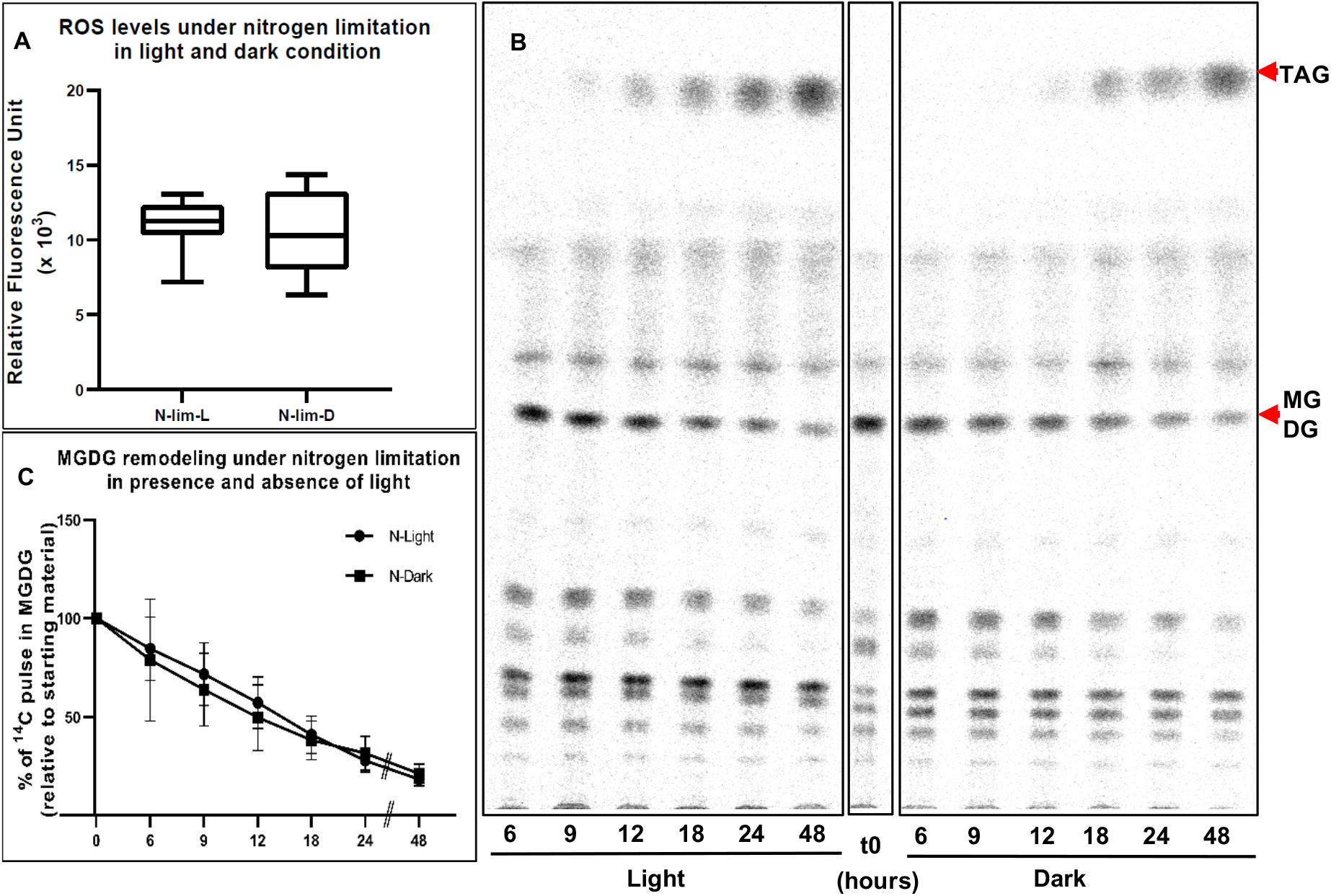
Photosynthesis as a source of ROS. A) ROS levels of N lim culture grown under light (N-lim-L) and dark (N-lim-D) regime as measured by DCFDA fluorescence after 24 hrs of inoculation. Fluorescence relative to O.D. B) A representative TLC-radiograph showing the remodeling of different lipids under nitrogen limitation in light or dark regime. C) Quantitative measurement of degradation of pre-formed MGDG relative to the starting levels of ^14^C pulse in MGDG in N-lim cultures grown in light and dark regime. Error bars indicate SD (n=3).

We next queried the transcriptome to find ROS generating enzymes that are differentially expressed under N-limitation. We found that expression of genes such as NADPH oxidase, xanthine oxidase/dehydrogenase, putrescine oxidase-like and copper amine oxidase (tyramine oxidase-like) are upregulated, as shown in Fig 7A. These enzymes all catalyze reactions that produce hydrogen peroxide or superoxide, depicted in Fig 7B. Roles of these enzymes in production of hydrogen peroxide in various other systems are well documented (Ma et al., 2016; Tavladoraki, Cona, & Angelini, 2016; Zandalinas & Mittler, 2018). Thus the altered gene expression we observe in ROS generating activities suggests that ROS accumulation is enhanced both by repression of ROS quenching activities and the induction of ROS generating activities.

**Fig 7.**
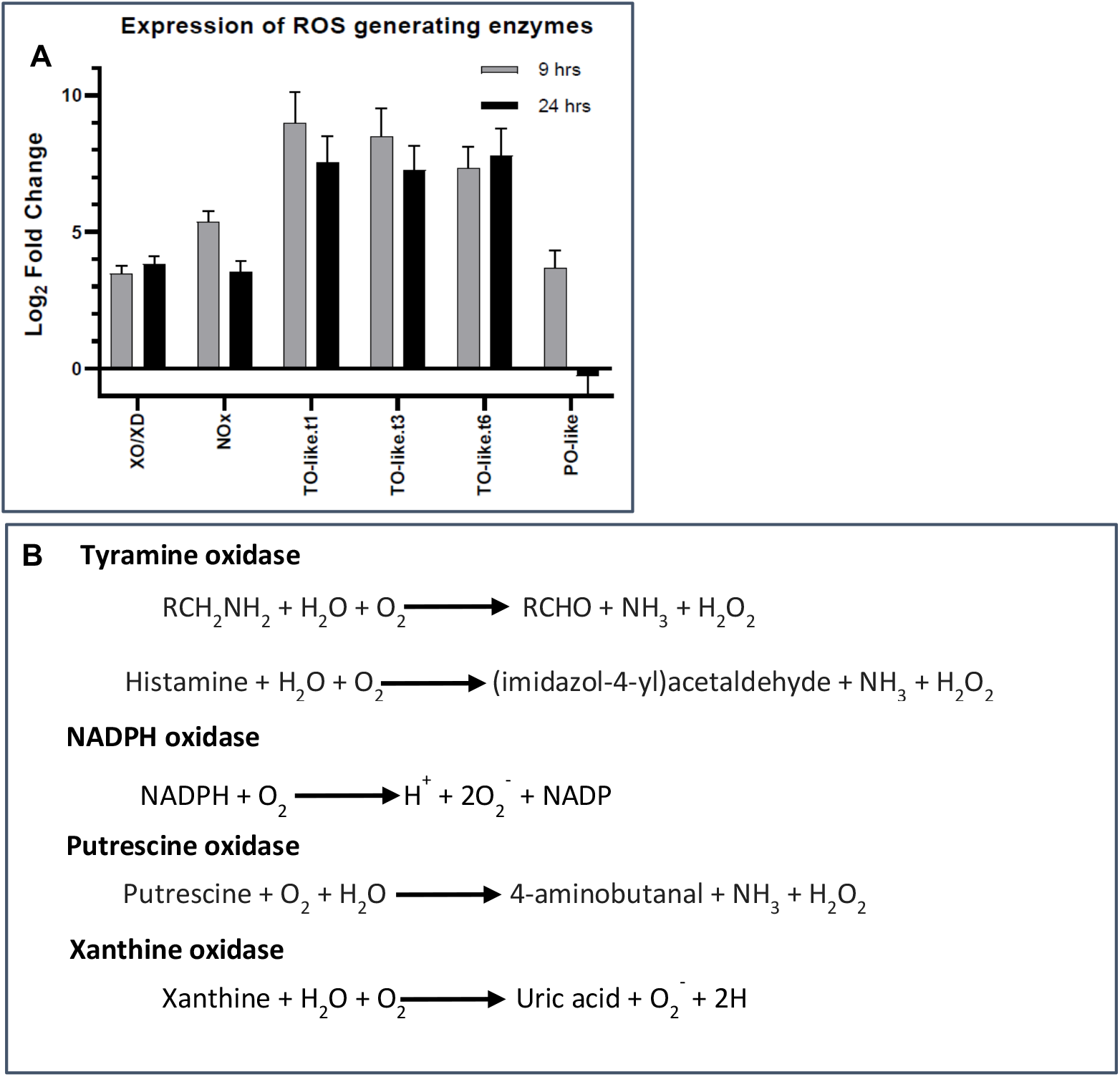
Sources of ROS. A) Expression of different ROS generating enzymes that increase during N limited growth as detected in the RNAseq studies. Error bars indicate SD (n=2). XO/XD, Xanthine oxidase/Dehydrogenase; NOx, NADPH Oxidase; TO-like, Tyramine oxidase-like; PO-like, Putrescine Oxidase-like. TO-like.t1, t3 and t6 indicate the three different splice variants encoded by the same gene. B) Enzymatic reactions catalyzed by these four enzymes showing the substrates and the products including the ROS. Source : KEGG pathway

### Role of NADPH oxidase in membrane remodeling

NADPH oxidases (NOX) are well studied enzymes given their various physiological roles in human health. Their functions are well researched in mammalian systems and are important drug target for various disorders (Meitzler et al. 2014). Therefore an array of chemicals are established as inhibitors of NOX (Altenhöfer et al., 2015). We used 2-Acetylphenothiazine, commonly known as ML171, to inhibit NADPH oxidase (Gianni et al. 2010). ML171 is a widely used inhibitor of NOX mediated ROS generation. When cultures were treated with 50 μM ML171 under N-limitation, we did not see any significant reduction in ROS level (Fig. 8A), as measured by DCFDA assay. Despite this, we observed a decrease in MGDG degradation when NOX activity was inhibited by ML171 (Fig. 8B). While there was no increase in N-limited growth upon ML171 treatment (Fig. 8C), we saw that accumulation of TAG was significantly reduced (Fig. 8D). These results suggest that a pool of superoxide produced by NADPH may be acting to mediate N-limited MGDG degradation and TAG accumulation.

**Fig 8.**
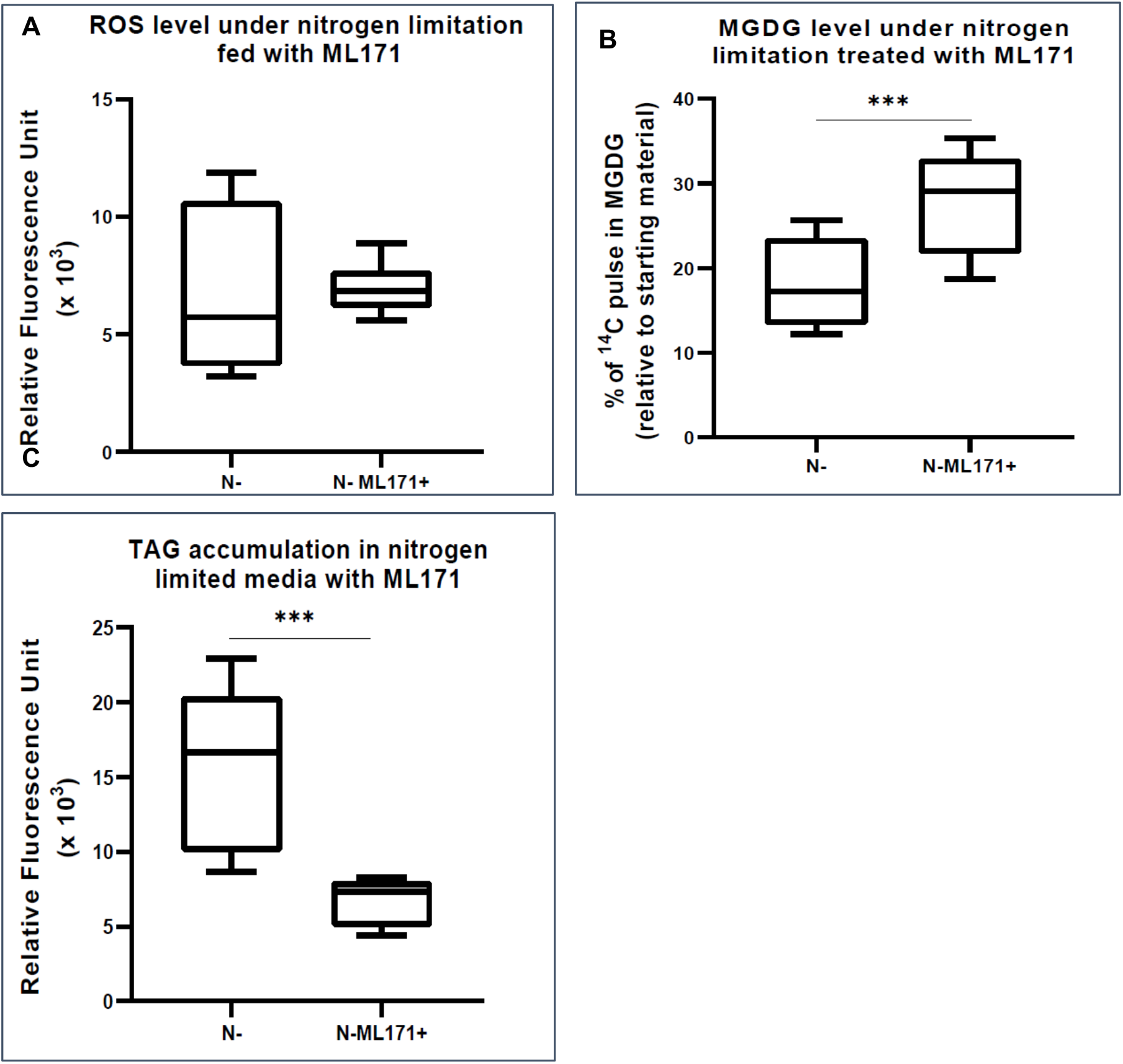
Role of NADPH oxidase in ROS generation and membrane remodeling. A) ROS level as measured by DCFDA fluorescence under N-lim condition when treated with NADPH oxidase inhibitor, ML171 (50 uM). ROS assay was carried out after 24 hrs of inoculation. Fluorescence is normalized to O.D. B) MGDG degradation under nitrogen limitation when fed with 50 uM ML171, (as measured after 24 hrs after inoculation). C) There is a substantial reduction in TAG level in cultures treated with ML171 when measured as nile red fluorescence at 24 hours post inoculation. All error bars indicate S.D (n=3). *** indicates a p-value <0.0001 in a paired student t-test.

**Fig 9.**
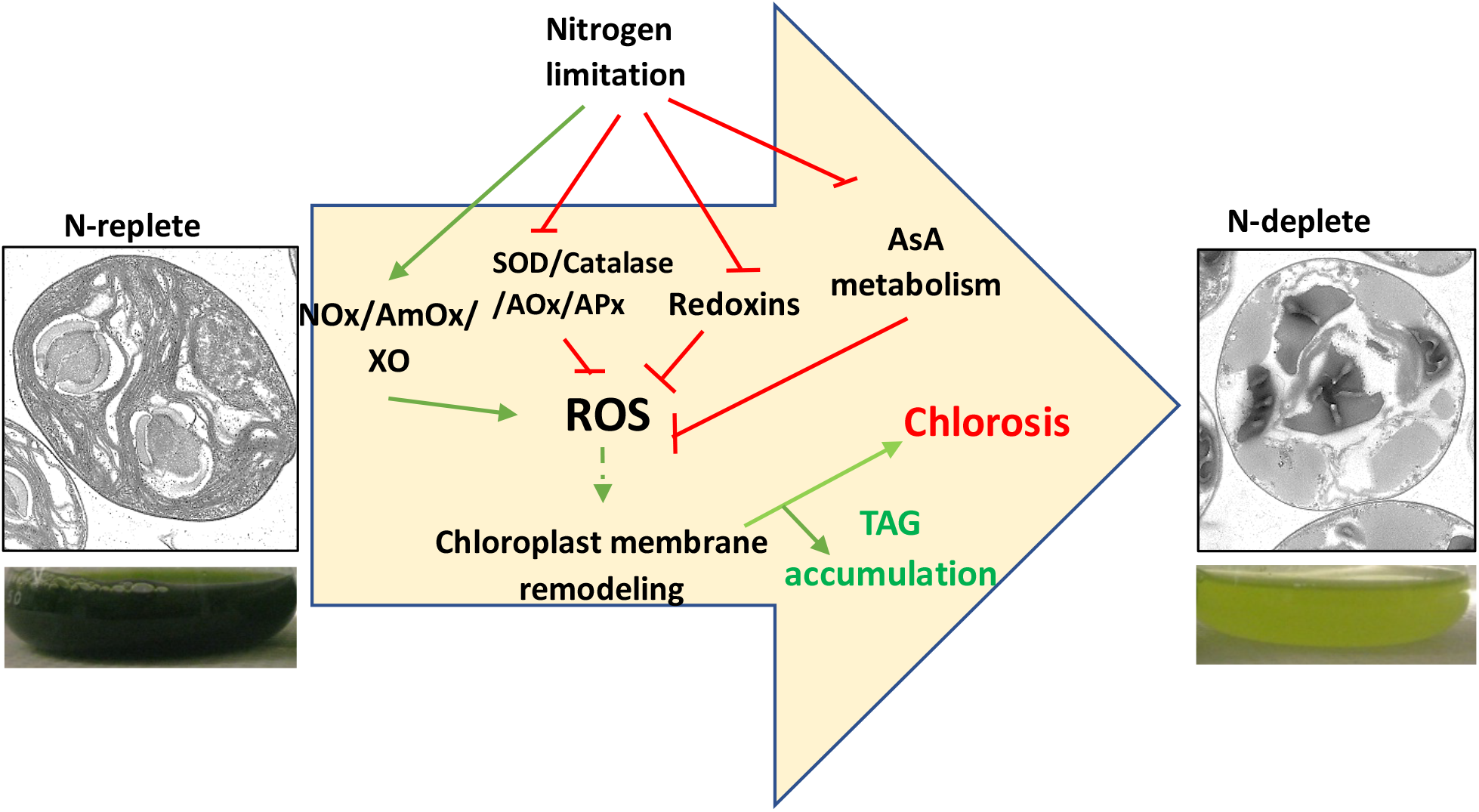
Proposed model for Nitrogen starvation mediated ROS accumulation leading to chlorosis and TAG accumulation. NOx: NADPH oxidase, AmOx: Amine Oxidase, XO: Xanthine oxidase, SOD: Superoxide Dismutase, AOx: Alternate Oxidase, APx: Ascorbate oxidase.

## DISCUSSION

Macronutrient limitation has been studied for decades as a potent inducer of storage compound (oil and starch) accumulation in oleaginous photosynthetic microorganisms, however the industrial usefulness of this mechanism of oil accumulation is invariably limited by a decline in photosynthetic capacity and biomass yield. Finding ways to stimulate oil accumulation without a penalty for growth or biomass accumulation has been pursued as a major goal of this area of study. Indeed a major driver of the US Department of Energy Aquatic Species Program (Sheehan, et al., 1998) was the identification of a hypothetical “lipid trigger” that could be induced under optimal growth conditions, leading to the reproducible and high-yield production of desired products. This supposed trigger has proved elusive, and the manipulation of single enzymes or transcriptional regulators has proven only marginally effective (Ajjawi et al. 2017). Another approach for the “lipid trigger” is the use of synthetic chemicals that induce storage lipid production with minimal retardation in growth has been shown recently, although the precise molecular mechanisms by which these compounds induce lipogenesis are, at present, poorly defined (Wase et al. 2015; 2014; 2019; Wase, Black, and DiRusso 2018). Despite this, significant progress has been made in understanding the enzymology and certain aspects of the regulation of TAG biosynthesis and degradation in algal model systems such as *Chlamydomonas* (Li-Beisson, Beisson, and Riekhof 2015; Li-Beisson et al. 2019) and *Nannochloropsis* (Ma et al., 2016), however the existence and nature of a hypothetical “lipid trigger” remains tenuous. This study presents several new insights related to signals of abiotic stress that are likely playing an underappreciated role in stress-induced TAG accumulation in oleaginous algae.

First, we showed that, as in other algal species, N-limitation in *C. sorokiniana* UTEX-1230 is a potent inducer of thylakoid membrane degradation and channeling of FA from chloroplast lipids into TAGs. Our transcriptomic analysis has revealed that many key enzymes that are involved in regulating ROS levels in cells, such as superoxide dismutase, catalase and stromal ascorbate peroxidase are downregulated, concomitant with an increase in ROS levels under N-limitation. This strongly suggests that cells are actively accumulating ROS in a regulated fashion under these conditions, by downregulating enzymes involved in quenching of ROS. The DCFDA assay provides a quantitative measurement of total ROS, leaving it unclear which specific species of ROS (superoxides, hydrogen peroxide, etc) is increased, and which are linked to membrane remodeling and TAG accumulation. It is also possible that each species has an independent role to play in the various branches of the N-starvation response. We are unable to comment with any certainty on the subcellular localization of these enzymes as *C. sorokiniana* is not yet amenable to genetic transformation. But by organelle preparation and biochemical assays, Kitayama *et al*., have shown that Fe-SOD was present in chloroplast and Mn-SOD was present in mitochondria of *Chlamydomonas reinhardtii* (Kitayama, et al., 1999). In the case of *C. sorokiniana*, we find that expression of both Fe-SOD (SOD2, 3) and Mn-SOD (SOD1) were downregulated. While we observe that the expression of SOD4, a Mn-SOD, is increased, the relevance of this increase in expression is unclear. Catalase is known to be present in mitochondria in *Chlamydomonas reinhardtii* (Kato, et al., 1997). Downregulation of ROS scavenging enzymes in both chloroplast and mitochondria indicates the possibility that more than one organelle could be a source of ROS under these conditions.

Expression of multiple ferredoxins and peroxiredoxins that are known to be involved in maintaining redox homeostasis are changed. Additionally we see that the expression of various ROS generating enzymes such as amine oxidases, xanthine oxidase and NADPH oxidases are increased. This indicates that *Chlorella sorokiniana* cells apply two pronged approach to ROS accumulation by downregulating scavenging systems and upregulating ROS producing processes, further indicating that ROS accumulation is an active and regulated process.

Data presented in this study pertaining to ROS accumulation and redox signaling are based on gene expression profiles, which may not always correlate with reduction of the function of the gene product, which can be dependent on the absolute levels of proteins and protein modification. However, we see a concerted reduction in the expression of many key players in ROS accumulation and redox signaling. These observations, coupled with an increase in ROS levels in the cell, indicates this is likely a transcriptionally regulated process, and that there is an underappreciated role for ROS and redox signaling under N-limitation. Increase in ROS levels under nitrogen limitation is also observed in the diatom *Phaeodactylum tricornutum* (Mizrachi et al. 2019). This indicates that accumulation of ROS under nitrogen limitation is not limited to *C. sorokiniana*, and possibly has a physiological advantage that has enabled this phenotype to be selected and maintained if these distantly related photosynthetic eukaryotes.

Ectopic induction of ROS by inhibition of SOD leads to MGDG degradation and quenching of ROS under nitrogen limitation reduces MGDG degradation. The concomitant reduction in TAG levels in ascorbate-treated N-limited cells is thus likely due to the reduction in degradation of membrane lipids such as MGDG and phospholipids. From these observations, it is clear that membrane remodeling mediated TAG accumulation is closely linked with redox signaling. In *C. reinhardtii*, significant flux of fatty acid for TAG accumulation is derived from preexisting glycerolipids (Young, & Shachar-Hill 2021). This is evident by the fact that mutations in genes encoding enzymes such as PGD1 and PDAT1 leads to reduction in TAG accumulation under N-limitation. PGD1 and PDAT1 are enzymes involved in turnover of fatty acid from MGDG and PC to TAG respectively (Li, et al., 2012; Yoon et al., 2012). Taken together, our study strongly implicates N-limitation induced ROS accumulation as playing a crucial role in the metabolic reprogramming that leads to TAG accumulation.. Future studies will be aimed at identifying the components that sense N-limitation and alter ROS accumulation and identifying the enzymes that act downstream of the ROS burst to regulate membrane turnover and TAG accumulation, thus providing additional details regarding the molecular trigger for macronutrient-limited TAG accumulation in microalgae.

## MATERIALS AND METHODS

### Culturing and growth measurements

*Chlorella sorokiniana* cells were grown in Tris Acetate Phosphate media (Gorman and Levine 1965) at 30°C under continuous light. For nitrogen limitation, parent cultures were grown for 2 days and then washed thrice in appropriate media before inoculating at a final concentration of 10^7^ cells/ml. Cells were inoculated into either control or nitrogen limited media with 10 mM or 250 μM ammonium chloride, respectively. Culture growth was measured as optical density (O.D) at 750 nm.

### Nile red staining for oil droplets

*C. sorokiniana* cells from appropriate treatment were first normalized to 0.2 O.D and then 10 μL of a 100 μg/ml stock of nile red (Sigma Aldrich) in DMSO was added to 100 μL of the culture. After 10 minutes of incubation the cells were used for fluorescence microscopy or measurement of fluorescence. Microscopy was carried using an EVOS-fl epifluorescence microscope with a GFP light cube. Quantitative fluorescence measurements were performed on a Synergy H1 hybrid reader with excitation and emission wavelengths of 455 and 560 nm respectively, and fluorescence values were normalized to the optical density.

### TEM analysis

For TEM analysis, cells were grown in appropriate media for four days and then fixed in formaldehyde before processing the samples. Sample preparation and analysis was carried out as described by Gojkovic et al. (2014.)

### Radiochemical labeling and analysis of lipids

Two days after inoculation, parent cultures were supplemented with ^14^C acetate (10^6^ cpm/ml of culture-specific activity of 56mCi/mMol-supplied by ARC Inc.,) for 8 hours. The cells were washed and inoculated in appropriate media at a concentration of 10^7^ cells/ml. 1 ml of culture was drawn out at each time point including the starting time point. Cell pellets were collected by centrifugation at 4000 rpm for 5 mins. Bligh and Dyer extraction was performed on the samples by partitioning lipids into the organic phase of a chloroform/methanol/0.2M KCl solvent system (1:1:0.8, v/v/v). Glycerolipids were dried under N_2_ and sequentially separated by TLC on a silica-G60 plate (EMD-Millipore) in two solvent systems. The first ascent to 2/3 the height of the plate in a solvent system of chloroform, methanol, acetic acid and water (85:12.5:12.5:3, v/v/v/v) was used to separate polar lipids. The plate was then dried and a second ascent to the full height of the plate was performed in petroleum ether, diethyl ether and acetic acid (80:20:1) to separate neutral lipids such as TAG, steryl-esters, and free sterols. After drying, plates were exposed to a storage phosphor screen for 24 hours and counts were read on a GE-Typhoon FLA 9500 scanner. Images were analyzed by ImageQuant TL v8.1, and individual bands were identified by reference to standards of known R_f_, and plotted as a percentage of the total counts for the lane.

### Transcriptome analysis by RNAseq

For RNAseq analysis we used two biological replicates with two technical replicates each of cultures grown in nitrogen replete and deplete media. Samples were prepared by pelleting the cells at 5000 rpm for 3 minutes and freezing them in liquid nitrogen. RNA was extracted by the trizol method. Briefly, frozen cells were gently suspended in 500 μL of Trizol reagent (Ambion TRIzol LS reagent) and incubated for 10 minutes at 4°C. 600 μL of chloroform was added, gently mixed and phase separated at 12,000 rpm for 10 minutes in 4°C. RNA was precipitated from the supernatant with 40 μL of NaCl (5M) and 500 μL isopropanol by incubating at 4°C for 10 mins. RNA was pelleted by centrifuging at 12,000 x g for 20 minutes at 4°C followed by washing the pellet with ethanol and further centrifugation for 3 minutes at 12000 rpm in 4°C. The pellet was carefully dried and suspended in RNase-free water. RNA library preparation and Illumina sequencing (75 bp single end read) was carried out by Genewiz (https://www.genewiz.com/). Reads were aligned to our in-house genome sequence of *C. sorokiniana* UTEX 1230 using Rsubread package (Liao, et al, 2013). Differential expression analysis was performed using the edgeR-limma workflow, and expression data for all genes are included as Supplementary Data File 1. (Ritchie et al., 2015; Robinson, McCarthy, & Smyth, 2009).

### ROS assay

Quantitative measurement of reactive oxygen species was carried out using DCFDA cellular ROS detection assay kit (Abcam, product number AB113851), and the assay was carried out following the manufacturer’s protocol. Briefly, the cells under different treatments were washed twice in the assay buffer and then normalized to 0.2 O.D and incubated with DCFDA at a final concentration of 10 μM for 1 hour. Following the incubation, the cells were washed twice with appropriate media to remove excess dye, incubated for 1 hour and fluorescence intensity was measured with excitation and emission wavelength of 485 and 535 nm, respectively. Values were plotted relative to O.D. For time course assay of ROS, cells were loaded with 10 μM DCFDA as mentioned earlier and set on a time course analysis of the fluorescence with the parameters on a Synergy H1 hybrid reader.

### Inhibitors and supplements

250 μM of diethyldithiocarbamate (Sigma Aldrich) in DMSO as a carrier was used to inhibit superoxide dismutase to induce the accumulation of ROS. 500 μM ascorbate (Sigma Aldrich) was supplemented to nitrogen limited culture to quench hydrogen peroxide. 50 μM ML171 (Selleck Chem) dissolved in DMSO was used to inhibit NADPH oxidase..

